# The solution path of the Li-Stephens haplotype copying model

**DOI:** 10.1101/2022.08.03.502674

**Authors:** Yifan Jin, Jonathan Terhorst

## Abstract

The Li-Stephens (LS) haplotype copying model forms the basis of a number of important statistical inference procedures in genetics. LS is a probabilistic generative model which supposes that a sampled chromosome is an imperfect mosaic of other chromosomes found in a population. In the frequentist setting which is the focus of this paper, the output of LS is a “copying path” through chromosome space. The behavior of LS depends crucially on two user-specified parameters, *θ* and *ρ*, which are respectively interpreted as the rates of mutation and recombination. However, because LS is not based on a realistic model of ancestry, the precise connection between these parameters and the biological phenomena they represent is unclear. Here, we offer an alternative perspective, which considers *θ* and *ρ* as tuning parameters, and seeks to understand their impact on the LS output. We derive an algorithm which, for a given dataset, efficiently partitions the (*θ, ρ*) plane into regions where the output of the algorithm is constant, thereby enumerating all possible solutions to the LS model at one go. We extend this approach to the “diploid LS” model commonly used for phasing. We demonstrate the usefulness of our method by studying the effects changing of *θ* and *ρ* when using LS for common bioinformatic tasks. Our findings indicate that using the conventional (i.e., population-scaled) values for *θ* and *ρ* produces near optimal results for imputation, but may systematically inflate switch error in the case of phasing diploid genotypes.

## 1 Introduction

Statistical analysis in genetics often requires evaluating the likelihood of a sample of genomes under a model of evolution. Unfortunately, this computation can rarely be performed exactly, because it requires integrating over the astronomical number of possible ancestry scenarios that could have generated the data. In 2003, Na Li and Matthew Stephens (Li and Stephens, 2003) proposed to approximate this intractable likelihood by modeling a newly sampled chromosome as a perturbed mosaic those previously observed. Simple yet effective, the Li-Stephens (LS) haplotype copying model has had a lasting impact in genetics and bioinformatics, with important applications to genotype imputation, phasing, linkage mapping, detecting natural selection, and other areas (Song, 2016).

LS depends on two parameters, *θ* and *ρ*, which are usually interpreted as the rates of mutation and recombination per unit time. Curiously, however, the model is not cognizant of time: in genealogical terms, it assumes that the sampled chromosome finds common ancestry with any other member of the population at a pre-determined number of generations in the past (Paul and Song, 2010). Since, in real data, there will be wide variation in the of age of ancestry at different locations in the genome, the interpretation of *Θ* and *ρ*, and their effect on inference, is not altogether clear—a fact which was already noted by Li and Stephens in their original paper.

In this study, we explore an alternative, non-biological perspective of *θ* and *ρ*, choosing to view them instead as tuning parameters in a machine learning algorithm. Then, the salient objective becomes understanding their effect on the output of the LS model. We derive a new, efficient algorithm for determining the complete solution surface of both the haploid and diploid variants of the LS algorithm. That is, for a given data set, the algorithm partitions the (*θ, ρ*) plane into regions such that the output of LS is constant within each region.

Our algorithm can be viewed as characterizing the trade-off between the effects of recombination and mutation: as the ratio *ρ/θ* tends to zero, recombinations become increasingly less likely, and the LS model simply copies from the most closely related haplotype in its entirety, at the potential expense of many mismatches. Conversely, as *ρ/θ* → ∞, there is free recombination between neighboring markers, and the LS model is able to find a path which is identical-by-state at every position (assuming no alleles are private to the focal haplotype), at the expense of improbably many recombinations. Our contribution is to characterize the behavior of LS for all intermediate values of *ρ/θ* as well, using an efficient procedure that requires only a single pass over the data.

For readers who are familiar with ℓ_1_-regularized regression (the LASSO), this can be seen as a type of LARS algorithm (Efron et al., 2004) for the LS model. Of course, the LASSO is regression, whereas the haplotype estimation problem addressed by LS is strictly unsupervised in practice. However, by applying our method to simulated data where the ground-truth ancestry is known, we can gain better insight into how the LS model functions, which can then be transferred to real-world applications.

The rest of the article is organized as follows. In Section 2, we first give a brief review of the LS model, and then derive our main results. Section 3 illustrates how these algorithms may be applied in practice to quantify the trade-off between recombination and mutation via simulation studies. Section 4 concludes with some discussion.

## 2 Methods

In this section we define the LS model, introduce our algorithms, and prove their correctness. We note once and for all that in this paper we focus squarely on the *frequentist* variant of LS, which returns a copying path (or pair of them, in the diploid algorithm) through haplotype space, basically by running the Viterbi decoding algorithm to obtain the maximum *a posteriori* (MAP) hidden state path through a hidden Markov model. Some other formulations of the LS model adopt a Bayesian perspective, where uncertainty in the unobserved copying path is modeled via a posterior distribution over hidden copying states. The techniques we introduce here are not applicable in the Bayesian setting, since they characterize the way in which the MAP path of the LS model changes as *θ* and *ρ* vary.

### 2.1 Notation and definitions

LS is used to decode positional ancestry of a “focal” chromosome consisting of *L* linked markers, using a panel of *N* “template” chromosomes. Each chromosome may be represented as a *haplotype*, that is a vector in 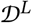, where 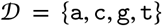 represent the four DNA nucleotides. The template haplotypes can be organized into a matrix

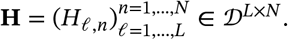

Throughout the paper, the variable 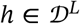 will be used to refer to a generic focal haplotype, and similarly the letter 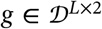 is used to denote a generic *diplotype*, that is a sequence of (unphased) diploid genotypes. We consider *h, g* and **H** as fixed instances of the above quantities, and will omit notational dependence on them when there is no chance of confusion.

For a positive integer *z*, the set {1,2,…, *z*} is denoted by [*z*]. A *path* (*of length ℓ*) is a sequence *π* = (*π*_1_,…, *π_ℓ_*) ∈ [N]^ℓ^ which characterizes the haplotype in **H** from which *h* copies at each position 1,…, *ℓ*. The notation |*π*| is used to denote the length of a path, so |*π*| = ℓ for a path of length *ℓ.*

Given a path *π*, the function

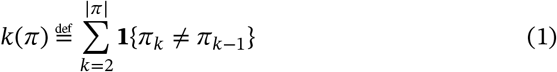

counts the number of times that *π* switches templates (i.e., the number of crossover recombinations). Similarly, the function

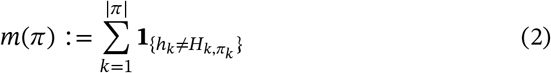

counts the number of mismatches between haplotype *h* and **H** for the copying path *π.* In the diploid case, if *π* and *λ* are two copying paths of equal length, then

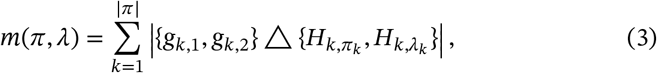

where *A* Δ *B* denotes the symmetric difference between sets *A* and *B*, is the number of panel mismatches for the focal diplotype *g.*

In Section 2.3 we will use some shorthand notation to refer to qualified subsets of the space of copying paths. A copying path *π* is an *ℓ*-path if |*π*| = ℓ. An Δpath for which *k*(*π*) = *r* is an (*ℓ, r*)-path, and similarly an (*ℓ, n*)-path is an *ℓ* path with the additional property that *π_ℓ_* = *n*. Lastly, an (*ℓ, r, n*)-path meets all three of these criteria.

### 2.2 The LS model

In its original formulation, LS is a generative model of the haplotype *h* conditional on the template set **H**. Formally, it is a hidden Markov model: at each position, *h* selects a particular template *π_ℓ_* ∈ [*N*] from **H**, whose identity is latent and unobservable. Conditional on this selection, the template allele is faithfully copied to h, except with some small error probability *p_θ_*. The “copying path” *π* ∈ |*N*|^*L*^ follows a stationary Markov chain: conditional on *π*_ℓ–1_, a switch occurs with probability *p_ρ_* ≪ 1/*N*; otherwise, with probability

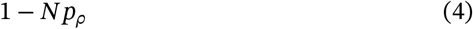

there is no switch and *π_ℓ_* = *π*_ℓ–1_. The leading factor *N* in (4) reflects the fact that, conditional on a switch having occurred between positions *ℓ* – 1 and *ℓ*, the identity of the newly selected haplotype at position *ℓ* is uniformly distributed among the *N* possible panel haplotypes. Similarly, the probability of correctly copying is 1 – 3*p_θ_*, where, again, the factor of 3 implies that the position mutates uniformly at random to one of the three other nucleotides not possessed by the template haplotype whenever a copying error occurs.

Thus, for a given *π*, the conditional likelihood of *h* is

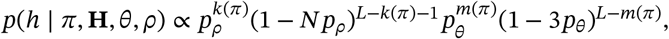

which leads to a compact expression for the negative log-likelihood (Lunter, 2019):

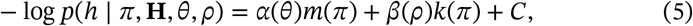

where *C* is a constant which does not depend on *π*, and we defined

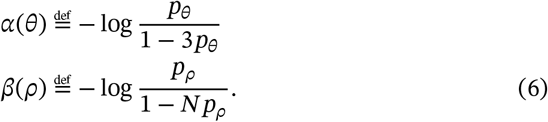

The function LS_*h*_(*θ, ρ*) is defined to return the lowest possible cost for (5):

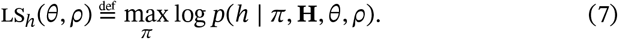

Li and Stephen’s original model is recovered by setting

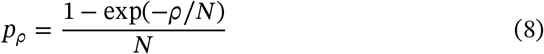

and 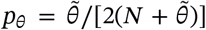, where the constant 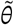 is derived by a population genetic argument (Li and Stephens, 2003, eq. A3). An alternative parameterization, based on a later, genealogical interpretation of LS (Paul and Song, 2010), is to set

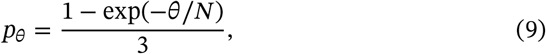

since the time to first coalescence between the focal and template haplotypes is roughly 1/*N* for large *N*.

An important difference between the original LS model and the one studied here is that, for reasons which become clear in the sequel, we assume that the probability of recombination is constant between each site. The same model was also recently considered by Lunter (2019), and is appropriate for large haplotype panels where the marker density is high and relatively uniform. It would not necessarily be appropriate for small data sets typed at a sparse set of markers.

Equation (5) asserts that log-likelihood of LS given *π* is, up to an irrelevant constant, simply a weighted combination of the number of template switches and sequence mismatches. Naturally, the weights depend on the mutation and recombination parameters, with higher values of *θ* (resp. *ρ*) leading to lower values of *α*(*θ*) (resp. *β*(*ρ*)), and correspondingly less weight placed on mismatches (resp. recombinations). Also, while it is technically possible for *α*(*θ*)or *β*(*ρ*) to be negative in (5), this requires very high rates of mutation and/or recombination which are not encountered in practice. In what follows, we assume min{*α*(*Θ*), *β*(*ρ*)} > 0. Note that this always holds if *p_θ_* and *p_ρ_* are set via (8) and (9).

### 2.3 Calculating all possible haploid decodings

In this section we derive an algorithm partition(*h*) to efficiently calculate all possible haploid decodings for various settings of *θ* and *ρ*. That is, for a given focal haplotype *h*, partition(*h*) returns a partition *S*_1_,…, *S_K_* such that

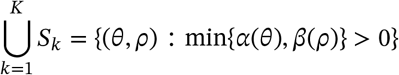

and for any *i* and (*θ, ρ*), (*θ′, ρ′*) ∈ *S_i_*,

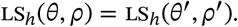

Note that there can be multiple *paths* that achieve the optimal cost LS_*h*_(*θ,ρ*); the regions returned by partition(*h*) have the property that the cost of any such path is the same within each region.

We arrive at the algorithm by a series of reductions. The first trivial result reminds us that, although LS is technically a two-parameter model, any choice of (*θ,ρ*) lies on a one-dimensional manifold of equivalent solutions.

#### Lemma 1.

*Let c* = *β*(*ρ*)/*α*(*θ*). *Thenforany θ‱, ρ′ such that*

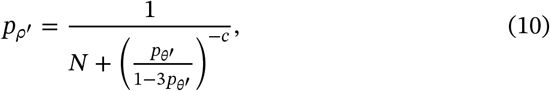

*we have LS*_*h*_(*θ‱, ρ′*) = LS_*h*_(*θ,ρ*).

*Proof.* If *ρ′* and *θ′* satisfy (10), then *β*(*ρ′*)/*α*(*θ′*) = *c*. Hence, by equation (5),

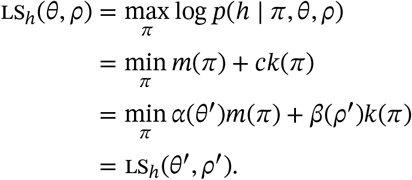

By the preceding result, we may assume that *α*(*θ*) = 1 in equation (5). Defining the optimal value function

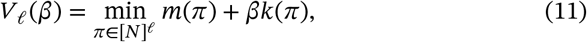

it therefore suffices enumerate the set {(*β, V_L_*(*β*)): *β* ≥ 0}. Next, define

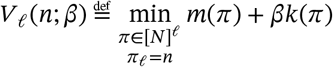

to be the optimal *ℓ*-path which copies from haplotype *n* ∈ [*N*] at the terminal position. Thus,

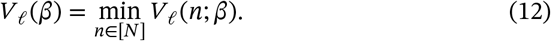

It follows from equations (1) and (2) that

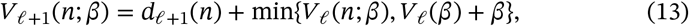

where *d_ℓ_*(*n*) is indicator function that whether there is a copying error from haplotype *n* at the terminal position *ℓ* + 1. It is easy to see that the functions *V_ℓ_*(*n*; *β*) and 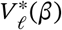 are piecewise linear and concave in *β.* Hence, dynamic programming can be used to solve (11) for all values of *β*, repeatedly applying (13) to determine the correct piecewise representation for *V_ℓ_*(*β*) and each *L.*

We experimented with this approach but found it to be too slow in practice. Equations (12) and (13) require taking the pointwise minimum of a collection of *N* piecewise linear functions. This entails finding all their points of intersection, which, though conceptually straightforward, is computationally burdensome for large *N.*

Instead, we derive an alternative algorithm that uses convex analysis to efficiently calculate *V_L_*(*β*). The algorithm recurses on a different quantity

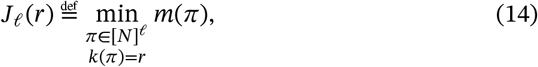

which is the least number of mismatches among all (*ℓ, r*)-paths. We then use a theorem from the changepoint detection literature to relate *V_L_*(*β*) and *J_L_*(*r*).

The theorem and ensuing discussion rely on the following basic results and definitions from convex analysis. A set *K* is *convex* if for all *x,y* ∈ *K*, the line 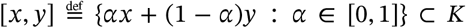. A point *x* ∈ *K* is a *vertex* if, for all *y,z* ∈ *K* such that *x* ∈ [*y, z*] (the line segment from *y* to *z*), either *x* = *y* or *x* = *z*. Given a set *X*, the *convex hull of X* is the intersection of all convex sets that contain *X.* If *X* ⊂ ℓ^2^ and |*X*| < ∞, the convex hull of *X* is a polygon, and can be completely described by the locations of its vertices. We use the notation conv(*X*) to denote the convex hull of a finite set *X* in the plane, and vtx(*X*) to denote the vertices of its convex hull.

The following key result is due to Lavielle (2005). We state it in an adapted form, and provide a short proof for completeness.

#### Theorem 2

(Lavielle, 2005). *Let*

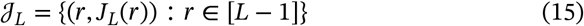

*be the graph of J_L_, and let r*_1_ < ⋯ < *r_M_ be such that*

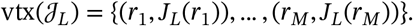

*Then*

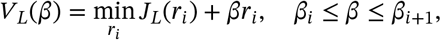

*where*

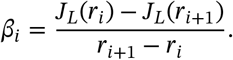

*Proof.* Since

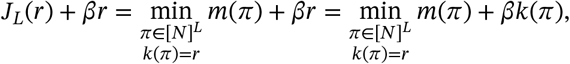

we have that

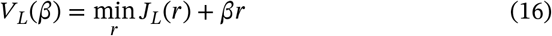

is the pointwise minimum of a collection of functions which are linear in *β*. Thus, there exists points *r*_1_ < ⋯ < *r_M_* such that *V_L_*(*β*) is piecewise linear, with vertices *β_i_* that satisfy

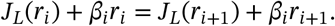

At each such *r_i_*, (16) implies that

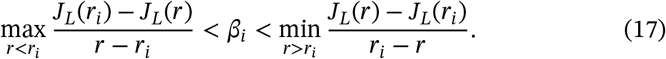

The preceding display establishes that (*r_i_, J_L_*(*r_i_*)) cannot be written as a convex combination of any two other points in 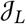, so it is a vertex of 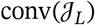.

By Theorem 2, determining *V_L_*(*β*) reduces to finding convex hull of the graph of *J_L_*(*r*). Now let

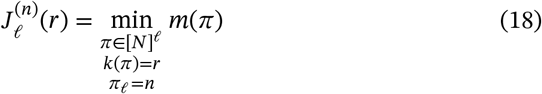

be the minimal number of mismatches among all (*ℓ, r, n*)-paths, and let

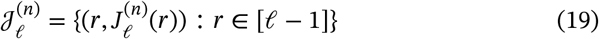

be its graph. We call an (*ℓ, r, n*)-path *π locally active* if 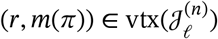. Similarly, an (*ℓ, r*)-path *π* is (*globally*) *active* if 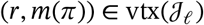.

By the preceding discussion, the set of active *L*-paths completely characterizes *V_L_*(*β*). The next result establishes that this set in turn may be obtained from the locally active *L*-paths. Let

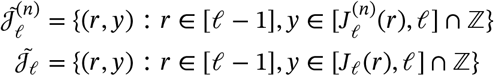

be the “truncated epigraphs” of *J_ℓ_* and 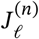, comprising all of the lattice points between the corresponding sets and the line *y* = *ℓ*. These sets have the same upper boundary and obviously (0, *ℓ*) and (ℓ −1, ℓ) are two common extreme points. Next, we characterize the extreme points of 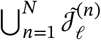:

#### Lemma 3.

*Let* 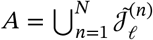 *and*

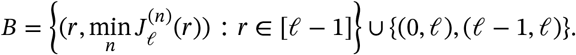

*Then* vtx(*A*) ⊂ *B*.

*Proof.* We have *B* ⊂ *A*, so let (*r,y*)∈ *A* \ *B*. Then either:

1. 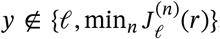, so that (*r,y*) can be written as the linear combination of (*r, ℓ*) and 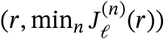; or
2. *y* = ℓ and *r* ∉ {0, ℓ – 1}, so that (*r,y*) is the linear combination of (0, ℓ) and (ℓ – 1, ℓ).

This shows that (*r, ℓ*) ∉ *B* ⇒ (*r, ℓ*) ∉ vtx(*A*), which is equivalent to the claim.

The following foundational result in convex analysis is stated for reference:

#### Theorem

(Krein-Milman). *If K* ⊂ ℝ^*d*^ *is compact and convex, then K* = conv(vtx(*K*)).

Since every set considered here is a finite set in ℝ^2^, the Krein-Milman theorem always applies.

#### Proposition 1.

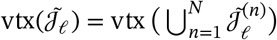.

*Proof.* Since 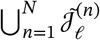 contains finitely many points, by the Krein-Milman theorem, its convex hull is spanned by its extreme points. Now by Lemma 3, the extreme points of 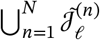 is a subset of

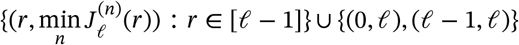

which is contained in 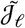 by definition. Thus, 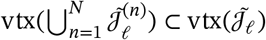. The other direction is by noticing 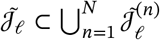.

At this point, we have reduced the original problem to that of finding the set of locally active (*L, n*) paths for *n* = 1,…,*N*. The next and final result shows how to compute these sets recursively. In theorem, we use an additional bit of notation: if *π* is an ℓ-path, and *c* ∈ [*N*], then we write *πc* to denote an “extension” (*ℓ* + 1)-path, such that (*πc*)_*i*_ = for *i* = 1,…, ℓ, and (*πc*)_ℓ+1_ = *c*.

#### Proposition 2.

*Let π = ϕn. If π is a locally active* (ℓ + 1, *r, n*)-*path, then either a*) *ϕ is a locally active* (*ℓ, r, n*)-*path, or b*) *ϕ is an active* (*ℓ,r* – 1)-*path*.

*Proof.* First suppose that *ϕ_ℓ_* = *n*. We claim that *ϕ* must be locally active. If not, then there exists a locally active (*ℓ, r*_1_, *n*) path *ϕ*^(1)^, a locally active (*ℓ, r*_2_, *n*) path *ϕ*^(2)^, and a number *α* ∈ (0,1), such that *r* = *αr*_1_ + (1 – *α*)*r*_2_ and

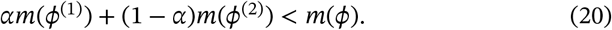

Adding *d*_ℓ+1,*n*_ = *αd*_ℓ+1,*n*_ + (1 – *α*)*d*_ℓ+1,*n*_ to both sides, we obtain

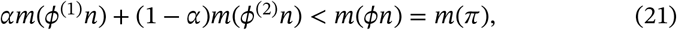

contradicting the fact that *π* is locally active.

Next, suppose that *ϕ_ℓ_* ≠ *n*. Then, since *π* is an (*ℓ* + 1, *r*)-path, *ϕ* is an (*ℓ, r* – 1)-path. If *ϕ* is not active, then one may similarly find active (*ℓ, r*_1_) and (*ℓ, r*_2_) paths *ϕ*^(1)^ and *ϕ*^(2)^ such that inequality (20) holds, where now *r* – 1 = *αr*_1_ + (1 – *α*)*r*_2_. Assuming without loss of generality that *r*_1_ < *r*_2_, this implies *r*_1_ < *r* – 1 < *r*_2_. Path *ϕ*_1_ may be extended to the (*ℓ* + 1, *r*_1_ +1, *n*)-path *ϕ*^(1)^*n*, and similarly for *ϕ*^(2)^, whence (21) holds. Because *r*_1_ + 1 < *r* < *r*_2_ + 1, we have *ϕ*^(i)^*n* ≠ *π* for *i* = 1,2. Thus, *π* is an interior point in the convex hull of all (ℓ + 1, *n*) paths, so it is not locally active. Hence, in either case we arrive at a contradiction.

**Algorithm 1.**
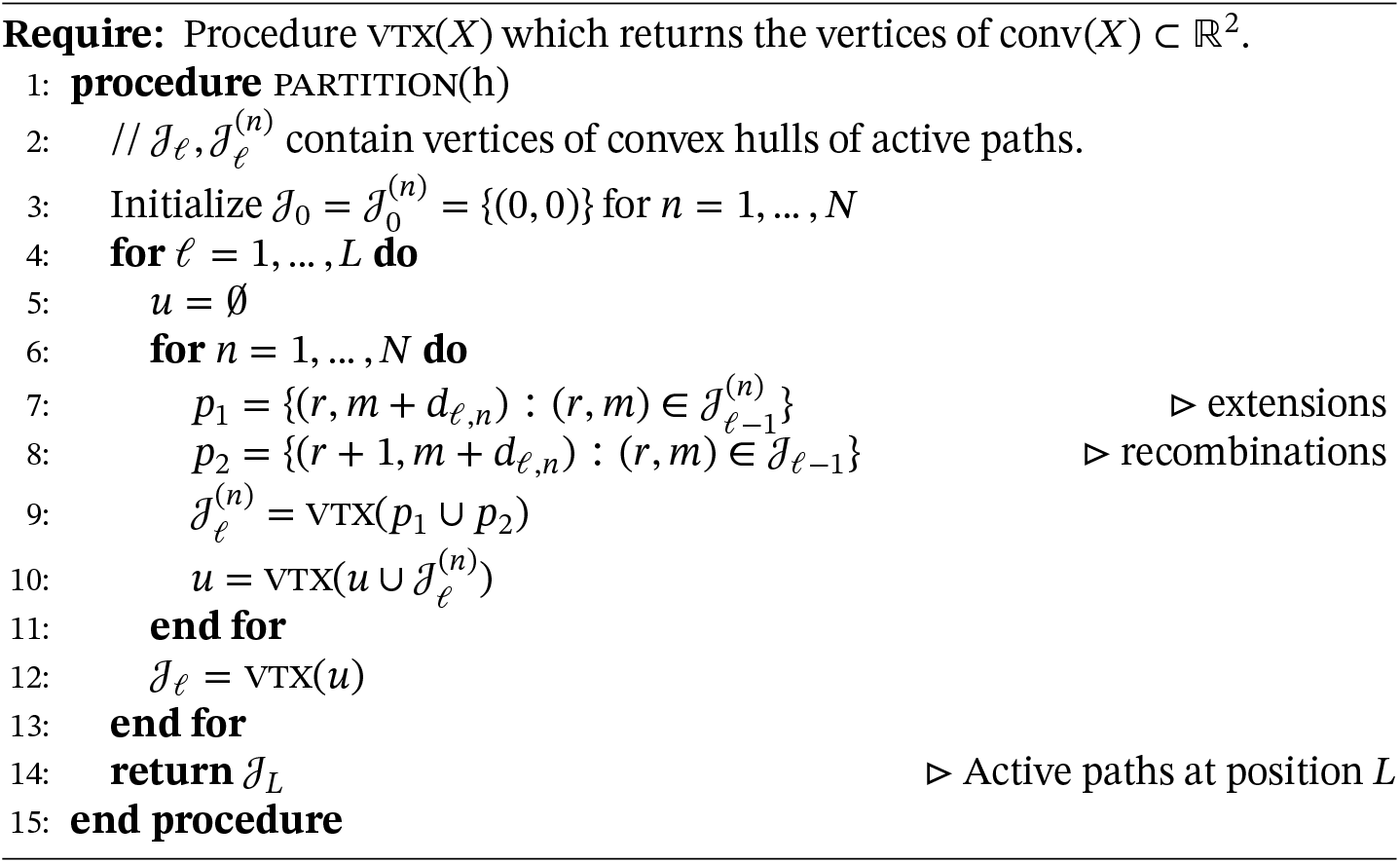
Haploid solution surface

By the preceding results, in order to find the set of active (*ℓ* + 1)-paths, it is only necessary to keep track of the set of active *ℓ*-paths, as well as the set of locally active (*ℓ, n*)-paths for each haplotype *n* ∈ [*N*]. Algorithm 1 implements Proposition 2. The output of the algorithm is 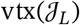. From this, Theorem 2 can be used to calculate ∞ = *β*_0_ > *β*_1_ > ⋯ such that LS_*h*_(1,*β*) is constant on each interval *β* ∈ (*β_i_, β*_*i*–1_). Finally, Lemma 1 and equations (8)–(9) yield the solution space for all (*θ,ρ*).

A few implementation details of Algorithm 1 are worth mentioning. As can be seen from lines 7–8, the assumption that *α*(*θ*) ≡ 1 causes the locally and globally active vertices to live on the lattice: 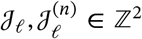. All numerical calculations are therefore exact, so the algorithm is impervious to rounding errors, or other floating point concerns. Also, for a finite set *X* ⊂ ℝ^2^, and assuming that the points in *X* are already sorted by their x-coordinates, the operation conv(*X*) used in lines 9 and 10 can be carried out in *O*(|*X*|) operations using e.g. Andrew’s algorithm (Andrew, 1979). This can easily be achieved by storing 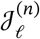 and 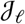 as sorted linked lists, and appropriately merging them in lines 9–10 instead of performing a naive set union. As the output of Andrews’ algorithm remains sorted, this ensures that the number of operations needed to perform lines 7–10 is minimized for all ℓ. It should be noted that, in practice, these optimizations may not improve performance unless *L* and *N* are very large. Finally, lines 7–9 are embarrassingly parallel and can be performed simultaneously using *N* different threads. However, the final reduction step (line 10) requires synchronization.

### 2.4 The diploid algorithm

The diploid extension to the Li-Stephens algorithm (e.g., Marchini et al., 2007; Loh et al., 2016) finds a pair of copying paths (*π*_1_, *π*_2_) ∈ [*N*]^2×*L*^ that maximizes the probability of observing a sequence of diploid genotypes 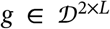. Similar to the haploid case, the log-likelihood of *g* given (*π*_1_, *π*_2_) has a compact expression (Lunter, 2019):

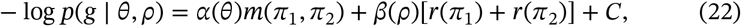

where *m*(*π*_1_, *π*_2_) was defined in equation 3.

We define LS_*g*_(*θ, ρ*) analogously to return a path pair 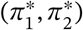 which minimizes equation (22). Clearly, Lemma 1 goes through without modification for LS_*g*_(*θ,ρ*) as well, so it is only necessary to determine the solution path for LS_*g*_(1,*ρ*). Algorithm 2 does this. The idea of the algorithm is similar to the haploid case, however more work is required in the form of an additional inner for loop needed to track both single and double recombination events. For each *n*_1_, *n*_2_ ∈ [*N*], the algorithm tracks a new set 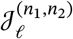 of locally active path *pairs*, as well as sets 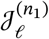 of “partially active” paths which lie on the convex hull of path costs involving haplotype *n*_1_ only. The set of “active” paths 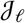 is now the convex hull of path costs taken over all possible path pairs.

The proof of correctness relies on a generalization of Proposition 2.

#### Proposition 3.

*Suppose that* (*πn*_1_, *λn*_2_) *is an active* (*ℓ* + 1, *r*, (*n*_1_, *n*_2_))-*path. Then one of the following is true*:

- (*π, λ*) *is a locally active* (*ℓ, r*, (*n*_1_, *n*_2_)) *path.*
- *π is a partially active* (*ℓ,r* – 1, *n*_1_) *path*.
- λ *is a partially active* (*ℓ, r* – 1, *n*_2_) *path*.
- (*π, λ*) *is an active* (*ℓ, r* – 2) *path*.

*Proof.* Similar to Proposition 2, the proof amounts to conditioning on last entries of *π* and *λ*, and showing that those paths must lie on the convex hull of the appropriate set of *ℓ*-paths. There are four cases to check depending on whether *π_ℓ_* = *n*_1_ and/or *λ_ℓ_* = *n*_2_. We prove one case and omit the repetitive details for the other three. Suppose that *π_ℓ_* = *n*_1_ and *λ_ℓ_* = *n*_2_, but that (*π,λ*) is not locally active. Then there are locally active (*ℓ, r*, (*n*_1_, *n*_2_)) paths (*ϕ*_1_, *ϕ*_2_) and (*γ*_1_, *γ*_2_) such that

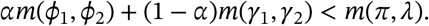

Thus

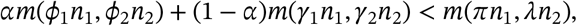

contradicting the supposition.

**Algorithm 2.**
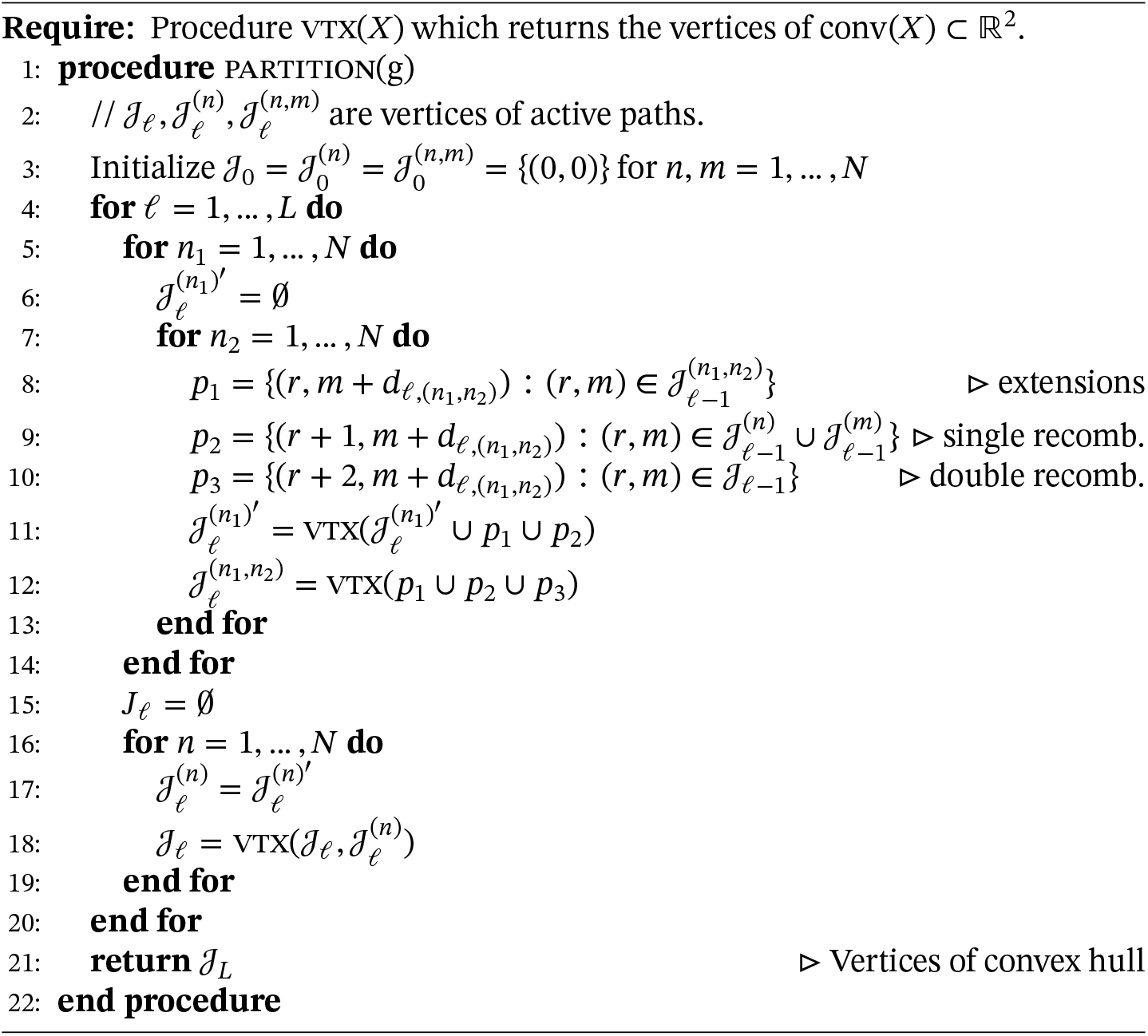
Diploid solution surface

## 3 Results

We used our algorithm to study two of the main use cases for the LS algorithm: phasing and imputation. The phasing problem attempts to resolve a sequence of diploid genotypes 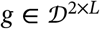 into a pair of haplotypes 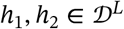, such that *g* and *h*_1_, *h*_2_ possess the same alleles at each position, and switching error is minimized. In the imputation problem, missing positions in a single haplotype are imputed using data from a reference panel. Notable phasing and imputation algorithms based on the LS model include fastPHASE (Scheet and Stephens, 2006), IMPUTE2 (Howie et al., 2009), MaCH (Li and Abecasis, 2006), SHAPEIT (Delaneau et al., 2012), and EAGLE (Loh et al., 2016).

### 3.1 Investigating imputation accuracy using Algorithm 1

For testing the Algorithm 1, we consider a haplotype imputation problem. Given a haplotype with the information of some SNPs is missing, we impute the haplotype with all possible *β* using the algorithm. To study imputation error, we considered the loss function

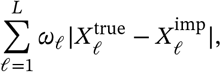

where the *ω_i_* are position-specific weights. We considered two choices for the weights: *ω_i_* ≡ 1, corresponding to Hamming distance between the imputed and true haplotypes; and *ω_ℓ_* = [MAF_ℓ_(1 – MAF_ℓ_)]^−1^, where MAF_ℓ_ is the minor allele frequency at position ℓ, thereby upweighting rare variants in the loss calculation.

The way we ran our algorithm is as follows: we first generate a focal haplotype *h* and reference panel *H.* The focal haplotype is then chosen as the underlying truth, and then all loci with minor allele frequency (MAF) less than 0.05 are discarded. We then use the retained loci to compute the solution surface, i.e. for a sequence of *β*, we compute the corresponding optimal path *p* = {*π*_*p*_1__,…, *π_p_k__*} with length *k* for each *β.* A missing locus is imputed by pasting the copying path state from the nearest flanking marker. The number of mismatches between the imputed copying path and the truth is computed in the end.

#### 3.1.1 Simulation study with fixed effective population size

We simulated 1001 sequences with 100Mb in a single population using msprime (Kelleher et al., 2016; Baumdicker et al., 2022). The length was chosen to be comparable to the size of a typical human chromosome. The effective population size was fixed to 1, and the scaled rates of recombination and mutation were both set to be 10^−4^ per unit of coalescent time. This resulted in a binary genotype matrix with roughly 300,000 rows and 1001 columns. For the haplotype imputation, we used the first column as the focal haplotype to be imputed, and the remaining columns 2–1001 as the haplotype panel. We then introduced missing data according to the MAF threshold mentioned above. This resulted in approximately 40% of the loci being retained on average.

For a given dataset, we first ran Algorithm 1 in order to find all possible LS paths. Then, for each interval of *β* where the LS solution has constant cost, we chose an optimal path and recorded its imputation error.^1^ Figure 1 shows the results of a single experiment for the Hamming loss. The curve is piecewise constant, with jumps at points where increasing *β* causes the cost of the optimal LS path to change. For this particular simulation, any setting 0.9 ≤ log(1 + *β*) ≤ 4.1 (roughly) was optimal in terms of imputation error. At the extremes *β* = 0 and *β* → ∞, we see the expected behavior. When *β* = 0 (i.e., *ρ* → ∞ in equation 6), there is free recombination so a copying path that contains zero copying errors (mutations) can be achieved. However, this results in some imputation errors since LD information is no longer being used for imputation. And as *β* → ∞, which implies *ρ* = 0 and complete linkage, the algorithm simply copies in entirety from the most closely related haplotype with no recombinations, resulting in many imputation errors.

**Figure 1:**
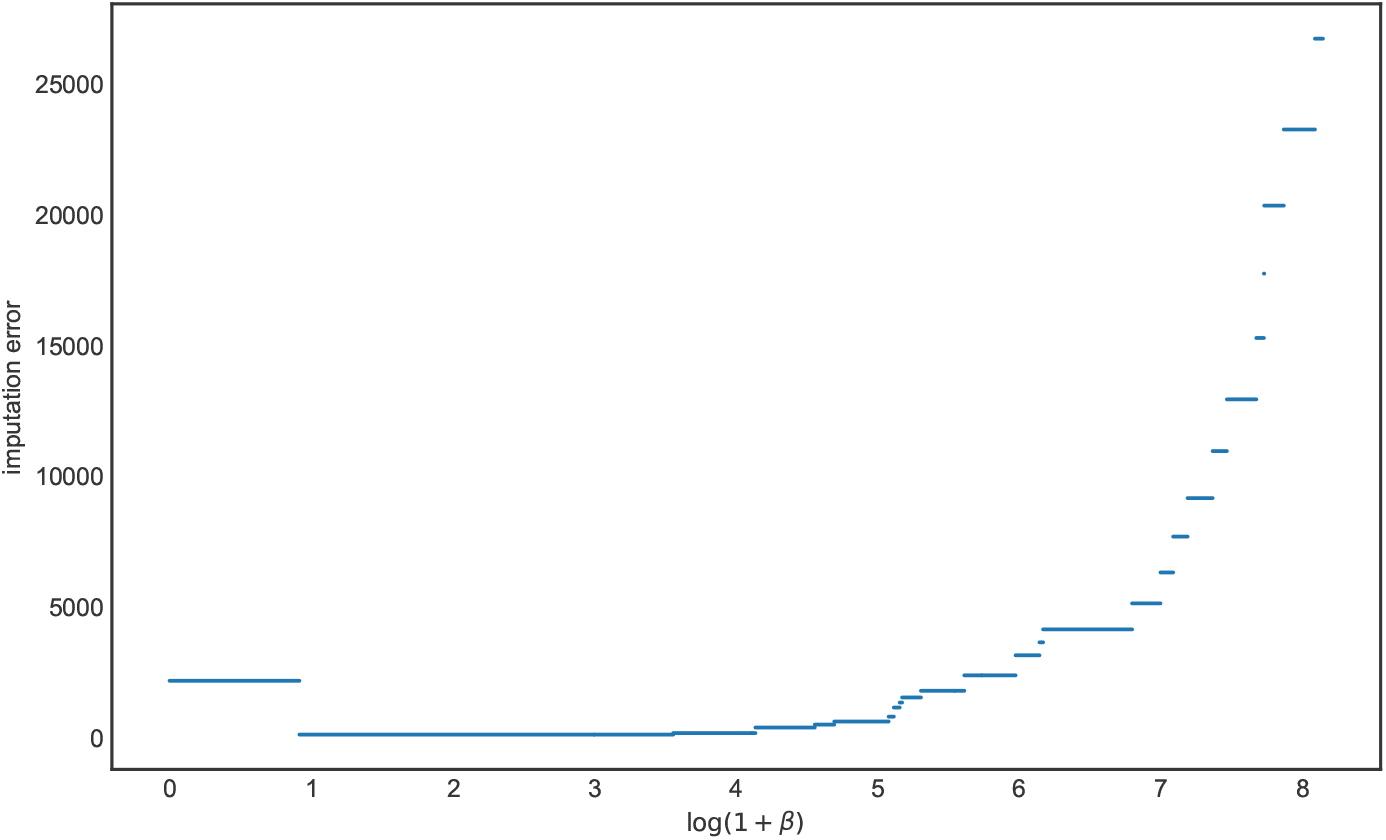
The imputation errors of all possible *β* for the haploid case, the *x* axis is the value of log(1 + *β)* and the *y* axis is the corresponding value of imputation error

We repeated this procedure 1,000 times and for each iteration, we determined the interval of *β* for which imputation error was minimized. Figure 2 depicts these results. Each box in the plot represents corresponds to an interval which was optimal in at least one run, with the height of the box representing the number of times it was the optimal interval across all 1,000 runs. (Because the corners of each box are all integers, they are displayed with transparency and a small amount of jitter to reduce overplotting.) The red dashed line in the plot corresponds to setting *β* according to equations (6) and (8), suitably transformed using Lemma 1, where *ρ* is the population-scaled rate of recombination. If the red line lies inside an interval, it means the LS model run with the population-scaled rate of recombination has the optimal imputation error. Otherwise, the results of imputation could be improved by choosing a different setting of *β*(*ρ*).

**Figure 2:**
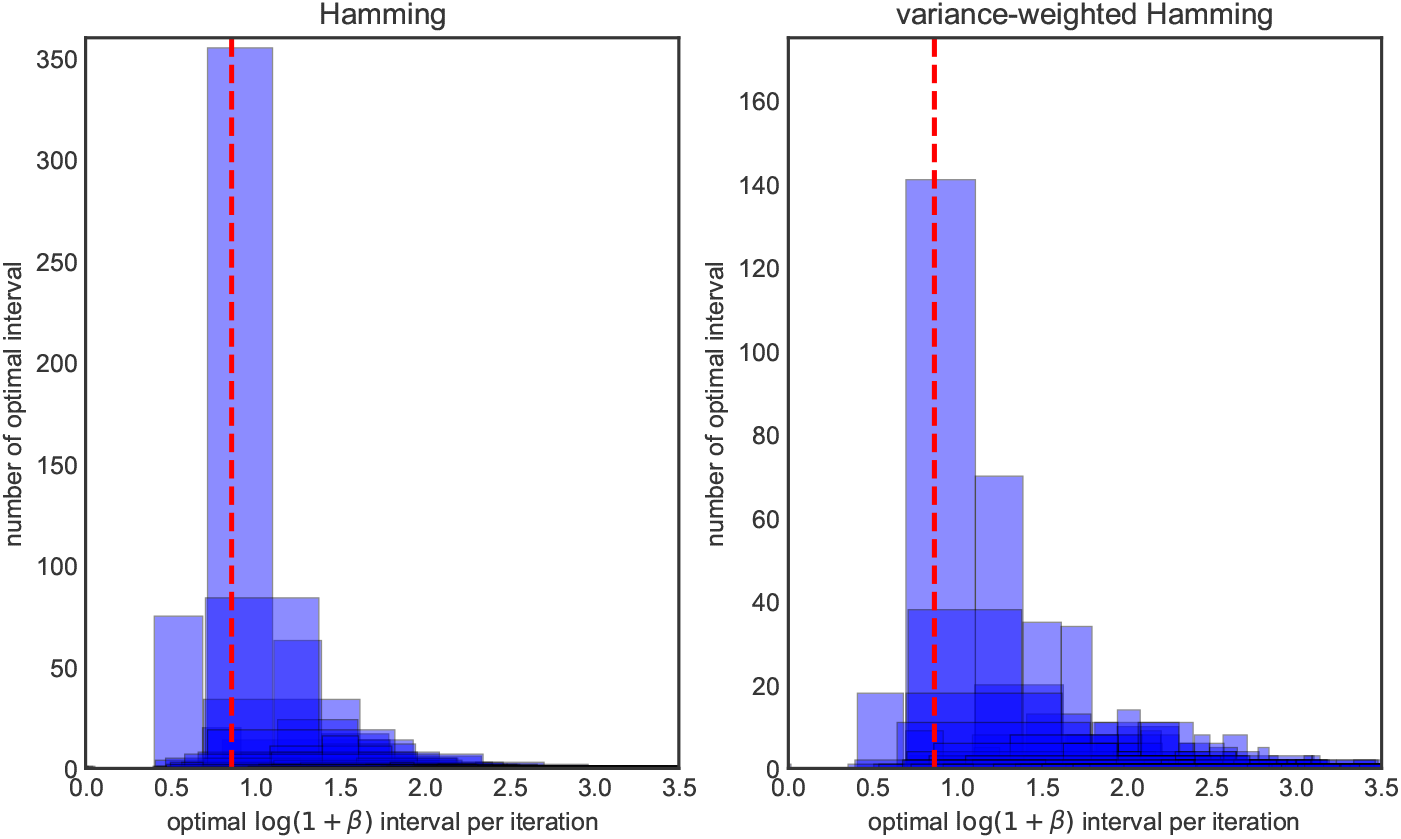
The histogram of optimal *β* intervals for algorithm 1, the *x* axis is the value of optimal log(1 + *β*) intervals in each iteration, the *y* axis is the number of replicates in 1000 iterations. The *x* axis of the red dash line is the true value of log(1 + *β*_0_) we used to generate data. The left panel is the histogram under the Hamming loss, the right panel is histogram under the weighted Hamming loss.

The left-hand panel of Figure 2 measures error in terms of Hamming loss, whereas the right-hand panel uses (inverse) variance-weighted loss. For Hamming loss, most of the optimal intervals are contained in log(1 + *β*) ∈ [0.5,2.0], and parameterizing LS using the population-scaled values for *θ* and *ρ* generally falls inside the optimal interval (in roughly 57% of the runs). The right-hand panel shows most of the optimal intervals lie are contained in between log(1 + *β*) ∈ [0.75,1.4). The same is mostly true for variance-weighted loss, however there is a longer right-tail to the distribution, and some simulations where setting *β* much larger (around 2.5-3) could lower imputation error. The percentage that the population-scaled values for *θ* and *ρ* falls inside the optimal interval is 0.246.

#### 3.1.2 Simulation study with variable effective population size

Next, we considered a more complex scenario where the population size varied according to a realistic model of human history. We simulated data using `stdpopsim` (Adrion et al., 2019), using the `Africa_1T12` demography for *H. sapiens.* This is a simplified two-population model with the European-American population being removed, and it describes the ancestral African population together with the out-of-Africa event (Tennessen et al., 2012). We simulated 1001 samples of human chromosome 2, but artificially reduced the length of the chromosome to be around 100Mb for computational reasons and to match the preceding experiment. The scaled mutation rate and recombination rate were around 8.7 × 10^-4^ and 9.8 × 10^-4^ respectively. We set the first sample as the focal and 2 to 1001 samples as the panel. The imputation procedure was the same as in the preceding section, where we retained the loci with MAF > 0.05 and imputed the remaining sites. To determine the population-scaled mutation rate, we set 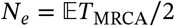, where 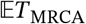 is the average time to coalescence in a sample of two chromosomes under the `Africa_1T12` demography, and then and then defined *θ* = 4*N_e_μ*.

For Hamming loss, Figure 3 shows the distribution of the optimal intervals is less dispersed than in the fixed population size case, with optimal log(1 + *β*) intervals concentrated between 0.75 and 1.2, which closely coincides with the population-scaled value (dashed red line). The percentage of runs where *β*(*ρ*) was optimal increased, to 0.73. Only a small amount of optimal intervals fall to the left of *β*(*ρ*). This indicates very occasionally, LS will perform better if the recombination rate is set lower than population-scaled value.

**Figure 3:**
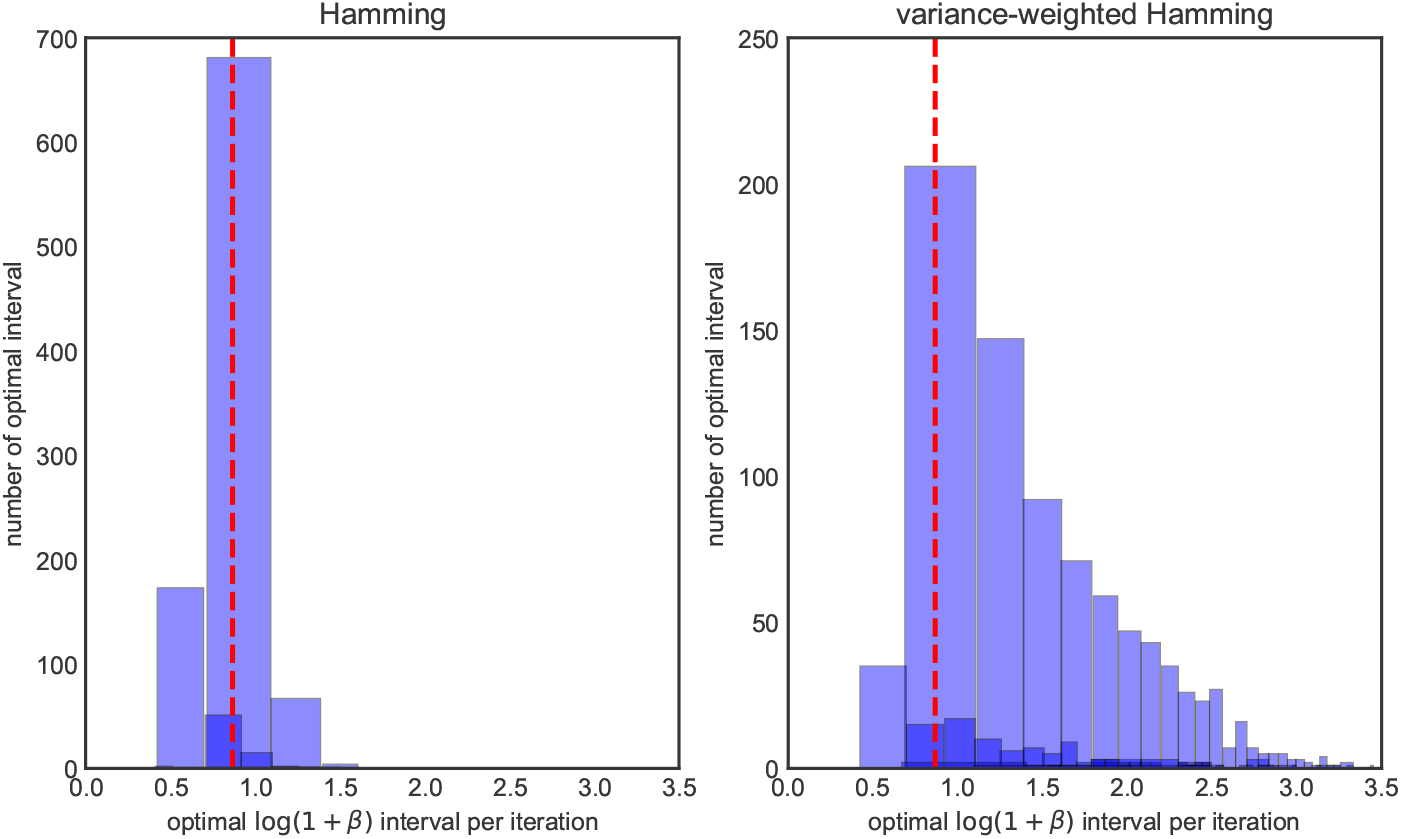
The histogram of optimal *β* intervals for algorithm 1 where the model is `Africa_1T12`, the *x* axis is the value of optimal log(1 + *β*) intervals in each iteration, the *y* axis is the number of replicates in 1000 iterations. The *x* axis of the red dash line is the true value of log(1 + *β*_0_) we used to generate data. The left panel is the histogram under the Hamming loss, the right panel is histogram under the weighted Hamming loss.

For the variance-weighted loss (Figure 3 right panel), we observed a similar phenomenon as in the constant-size case: there is a heavier right tail, and in a larger fraction of the simulations, imputation results could have been improved by setting *β* higher than the population-scaled value. The percentage of runs where *β*(*ρ*) was optimal decreased, to 0.223. However, in general, the previous two experiments show that the population-scaled rates should generally be adequate for phasing using the haploid LS algorithm.

#### 3.1.3 Accuracy of pre-phased imputation

A common workflow for imputing diploid genotypes is to first phase them into haplotypes and then run haplotype imputation (Howie et al., 2012). We repeated the preceding experiments to study diploid imputation using pre-phasing (Supplemental Figures A1–A4) and observed generally comparable results: the population-scaled values generally result in optimal performance for pre-phasing when considering Hamming loss, but there is a longer right tail when rare variants are upweighted in the loss calculation. For the more realistic, out-of-Africa demography, we observed less dispersion of the optimal intervals than for the constant demography.

### 3.2 Investigating phasing accuracy using Algorithm 2

For testing the algorithm 2, we considered a genotype phasing problem. Given a genotype sequence which is the pairwise sum of two haplotype sequences, and a reference panel, we aim to recover the information of each haplotype sequence. We first generated two focal haplotype sequences *h*_1_, *h*_2_ and reference panel *H* and combine *h*_1_ and *h*_2_ to form a genotype sequence. In order to get the phased haplotype sequences, we then fed the genotype sequence and reference panel to algorithm 2. We then measured the accuracy by measuring the switch error between the true and estimated haplotype sequences. Switch error was computed using the `--diff-switch-error` option of `vcftools` (Danecek et al., 2011).

#### 3.2.1 Simulation study with fixed effective population size

We again start with the simple simulation scenario where the effective population size is fixed and equal to 1. The value of mutation and recombination rate per time are both set to be 10^−4^. As noted above, the diploid solution surface algorithm scales quadratically in the reference panel size, as opposed to linearly for the haploid algorithm. For this reason, we considered a shorter sequence length and a smaller panel size. We set the sequence length to be about 10MB and use a reference panel 100 haplotypes. After generating 102 samples from the model, we choose the first two columns of the genotype matrix as the focal and 3 to 102 columns as the panel. An example of one run of the experiment is shown in Figure 4. In this experiment, the algorithm achieves minimum switch error when log(1 + *β*) is around 2.

**Figure 4:**
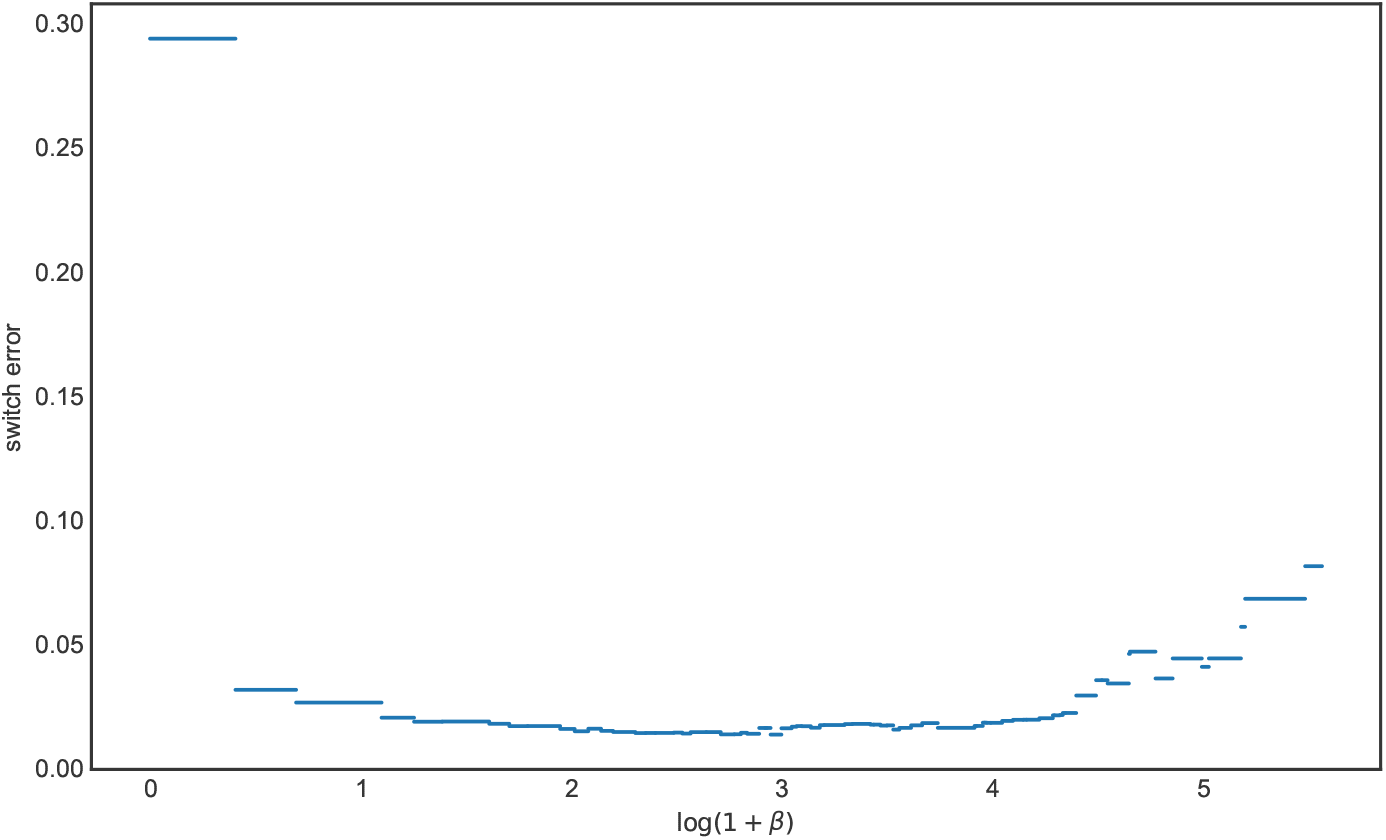
The switch errors of all possible *β* for the diploid case, the *x* axis is the value of log(1 + *β*) and the *y* axis is the corresponding value of the switch error

Figure 5 shows the results of running this experiment 1,000 times. In contrast to the haploid case, the optimal setting of *β* is systematically higher compared to the value based on the population-scaled rate, which is again shown as a red dashed line. Although the best *β* intervals seem to be more dispersed than the ones in the imputation problem, most log(1 + *β*) intervals are clustered on the right of the red dash line and are between 2 and 4. Moreover, only a few*β*s fall into the same partition as the red line. We also noticed for that some iterations, the optimal *β* interval is near 0. This can occur if there is one or more very closely related haplotypes in the reference panel. To validate these results, we compared the distribution of switch error using a *β* from the modal interval in Figure 5 (we chose 23.0), and compared it to that obtained when the *β* was suboptimally set according to the scaled rate of recombination. Figure 6 shows there is a difference of switch errors by using these two *β*s, with the distribution of switch error under the optimal setting possessing more mass at zero.

**Figure 5:**
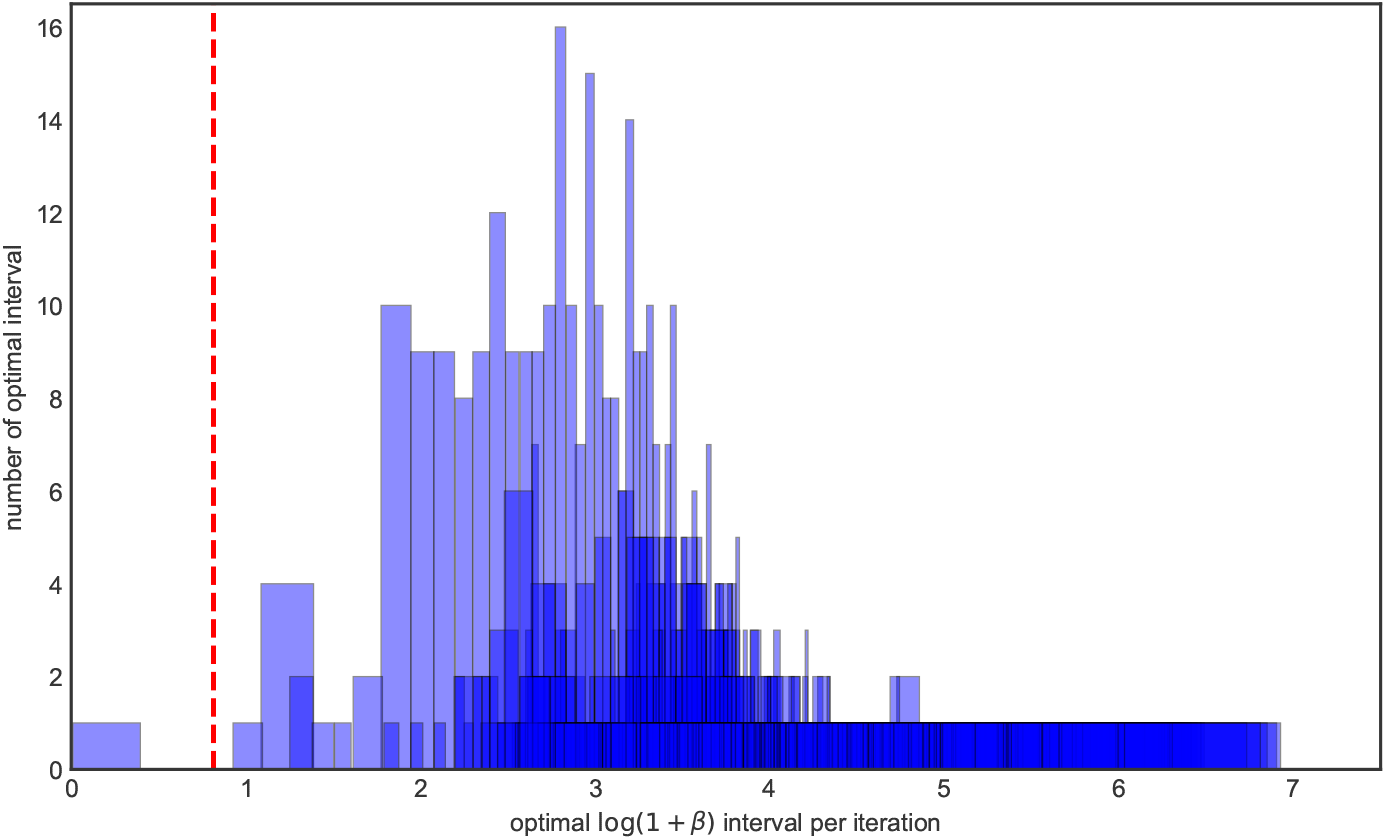
The histogram of optimal *β* intervals for algorithm 2, the *x* axis is the value of optimal log(1 + *β*) intervals in each iteration, the *y* axis is the number of replicates in 1000 iterations. The *x* axis of the red dash line is the true value of log(1 + *β*_0_) we used to generate data.

**Figure 6:**
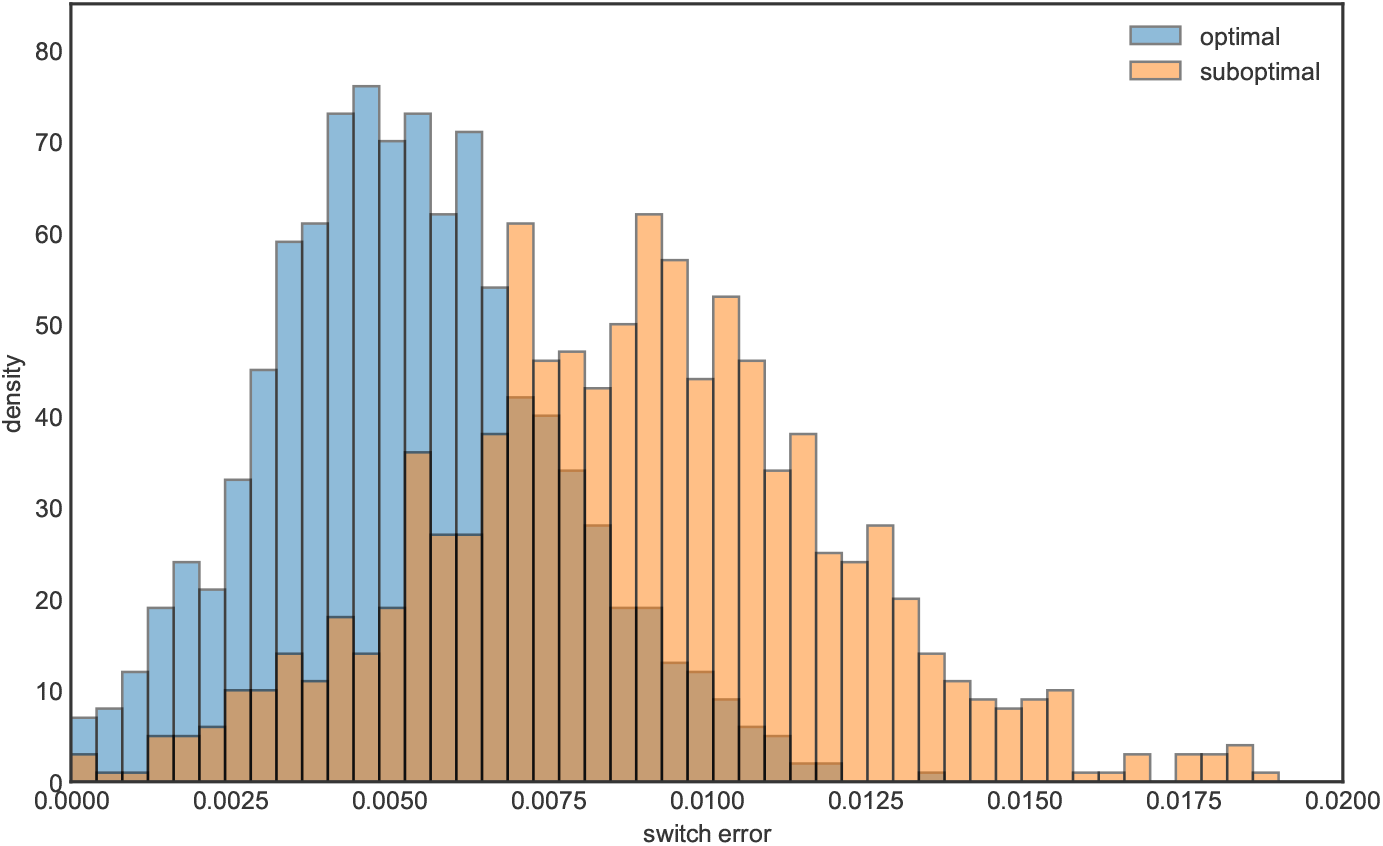
The histogram of switch errors for optimal and suboptimal *β*s respectively, the *x* axis is the value of switch error, the *y* axis is the density.

#### 3.2.2 Simulation study with variable effective population size

Finally, we repeated the phasing experiment using the `Africa_1T12` demographic model. We simulated 20% of chromosome 22, so that the sequence length is about 10Mb, and imputed a diploid genotype sequence using a reference panel of size 100. From Figure 7 we conclude the distribution of optimal log(1 + *β)* intervals has a similar pattern as in the fixed population size case. Increasing *β—*that is, penalizing recombinations more heavily—leads to lower switch error. Figure 8 shows the differences if we use optimal and suboptimal *β* respectively. The differences are more pronounced compared to the preceding section: more than half of the simulations using the “optimal” setting had lower switch error than almost every simulation using the “suboptimal” setting.

**Figure 7:**
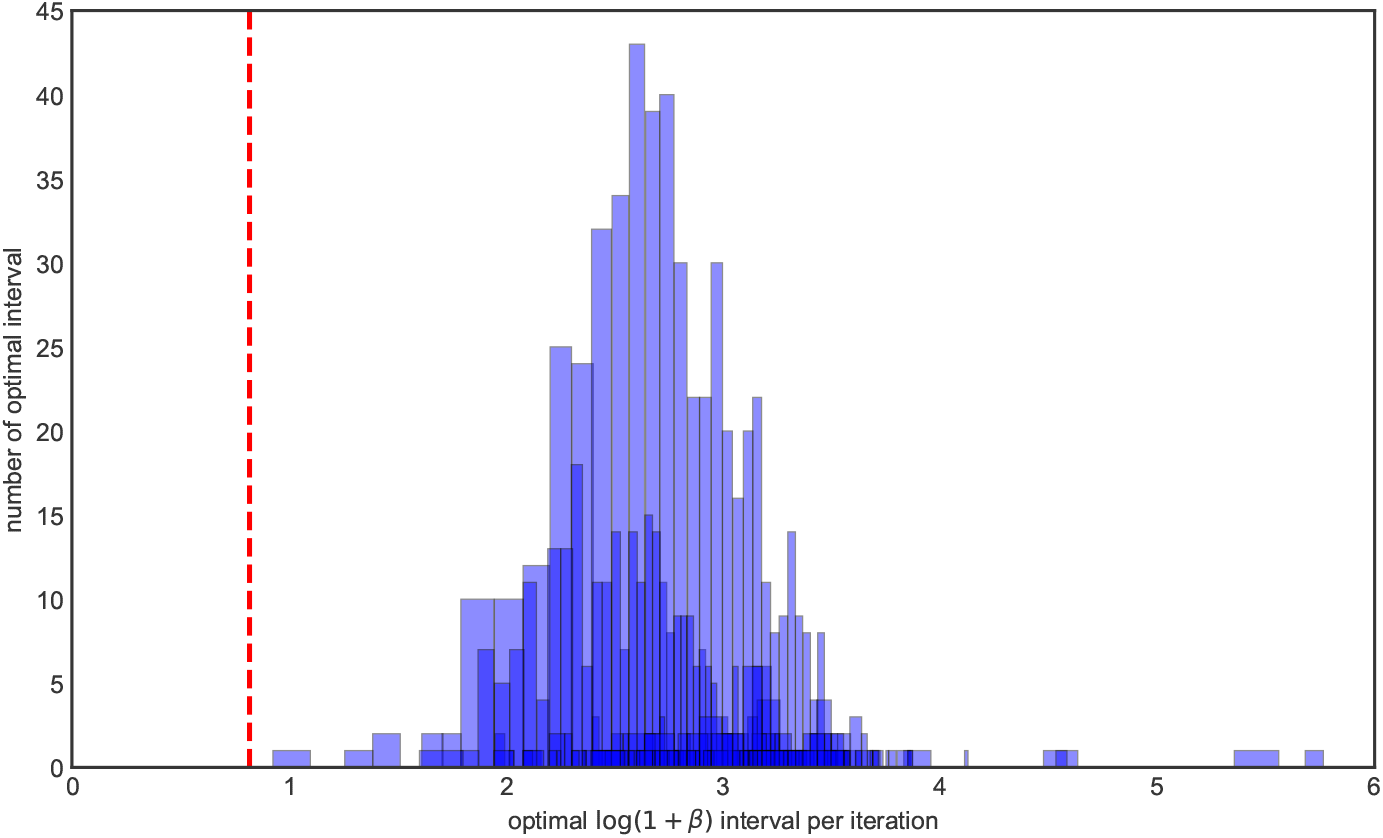
The histogram of optimal *β* intervals for algorithm 2 where the model is `Africa_1T12`, the *x* axis is the value of optimal log(1+*β*) intervals in each iteration, and the *y* axis is the number of replicates in 1000 iterations. The *x* axis of the red dash line is the value of log(1 + *β*_0_) where *β*_0_> is the truth.

**Figure 8:**
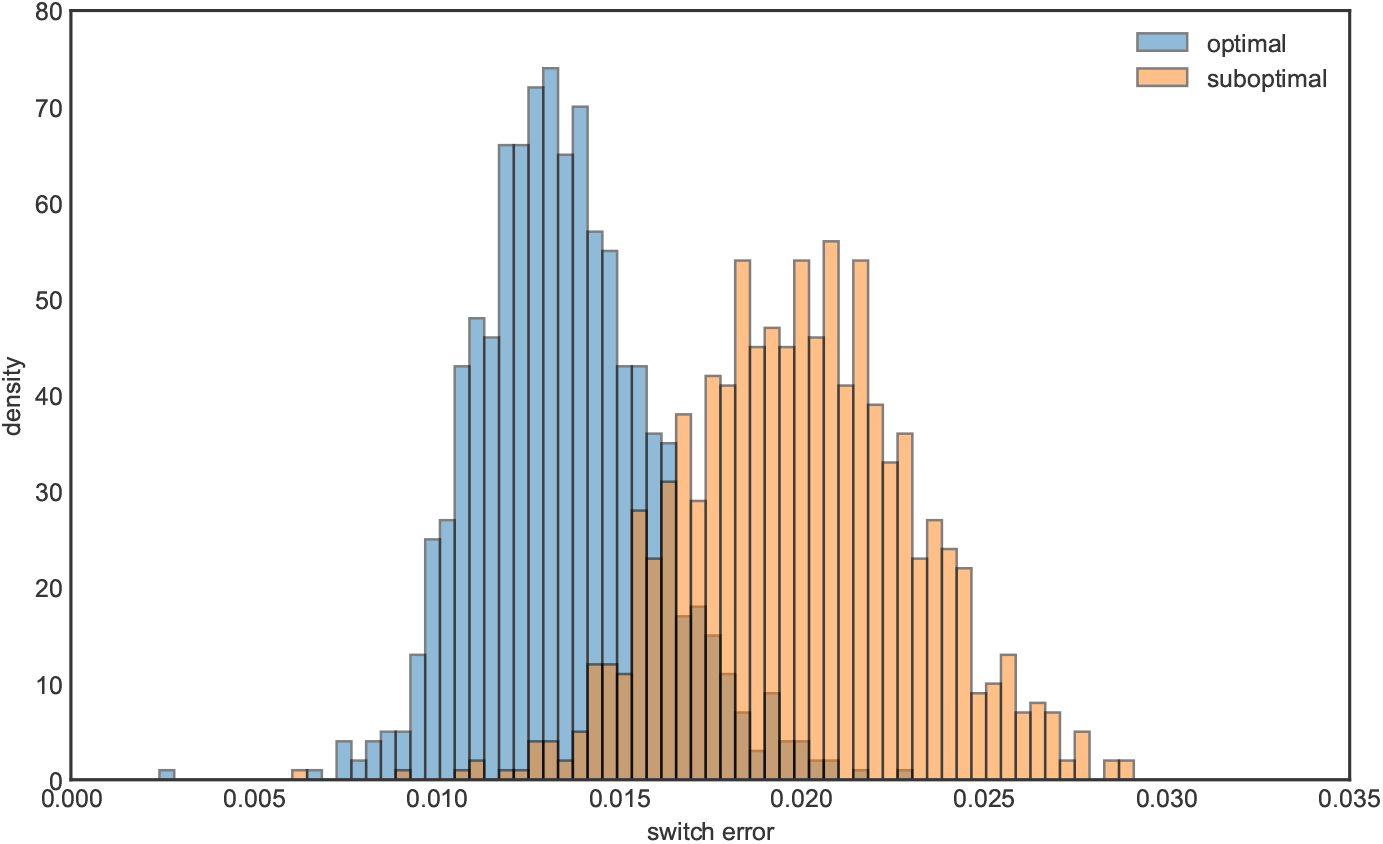
The histogram of switch errors for optimal and suboptimal *β*s respectively under the `Africa_1T12` model, the *x* axis is the value of switch error, the *y* axis is the density.

## 4 Discussion

In this paper, we derived a new algorithm for computing all possible solutions to the Li-Stephens haplotype copying model, as well as its diploid extension, as a function of the recombination rate parameter. Our results work by exploiting convex structure in the Viterbi decoding algorithm used to compute the optimal (frequentist) LS haplotype copying path. Our algorithms partition the LS parameter space into regions where the output of the model is constant. We showed how these can be useful for studying imputation and phasing accuracy, two of the most important uses of the LS model.

Our methods work by interpreting the LS model as a method for performing changepoint detection. Although this perspective appears to be new as far as the LS model goes (but see Ki and Terhorst, 2020), it has appeared in the literature before in other forms. The CROPS algorithm (Haynes et al., 2017) is a general procedure for computing the solution space of changepoint models as a function of a penalty parameter, which could also be applied here. The main difference between our contribution and theirs is that the CROPS algorithm is iterative, requiring multiple runs of the model in order to compute the entire solution surface, whereas our algorithm requires only a single pass over the data. Figures 1 and 4 illustrate that, for investigating derived quantities such as phasing or imputation error, it seems necessary to compute the entire solution surface, since the error curves do not posses any sort of regularity (e.g., convexity) which would allow one to know when a globally optimal solution has been found. However, for very large data sets, the iterative approach of the CROPS algorithm may be preferable.

Some further refinements to our algorithms are possible. While we showed in simulations that for diploid phasing that there is a gap between the *β* used for generating the samples and the optimal *β* for LS models, our computation of *β* is based on constant recombination rate. In contrast, most popular phasing and imputation packages, for example BEAGLE (Browning et al., 2021) or IMPUTE2 (Howie et al., 2009), use a recombination map whose value changes based on the genetic distance between each marker site. We do not see an easy way to modify our algorithm to accommodate this type of analysis since it is not even clear what the resulting output would be. A similar difficulty was noted by Lunter (2019) in the context of the fastLS algorithm.

The size of the reference panel considered in our simulation study is relatively small, especially for the diploid phenotype phasing setting, where we only used a reference panel with a size equal to 100. We had to choose this small value because the complexity of our Algorithm 2 is (at least) quadratic with the size of the panel. In contemporary imputation and phasing studies, the panel size is much larger, e.g. 2 × 10^5^ individuals in Browning and Browning (2016). Their study results indicate the imputation of low-frequency variants can be highly benefited from a large reference panel with accurately phased genotypes. A potential direction is thus to scale our algorithm to the setting where the size of the reference panel is large.

## Data and code availability

A software implementation of our method, as well as Jupyter notebooks which reproduce our results, are available at https://github.com/jthlab/lsss.

## Acknowledgements

We thank Hyun Min Kang for feedback on a draft of this manuscript. This research was partially supported by the National Science Foundation (grant number DMS-2052653).

## A Additional Figures

**Figure A1:**
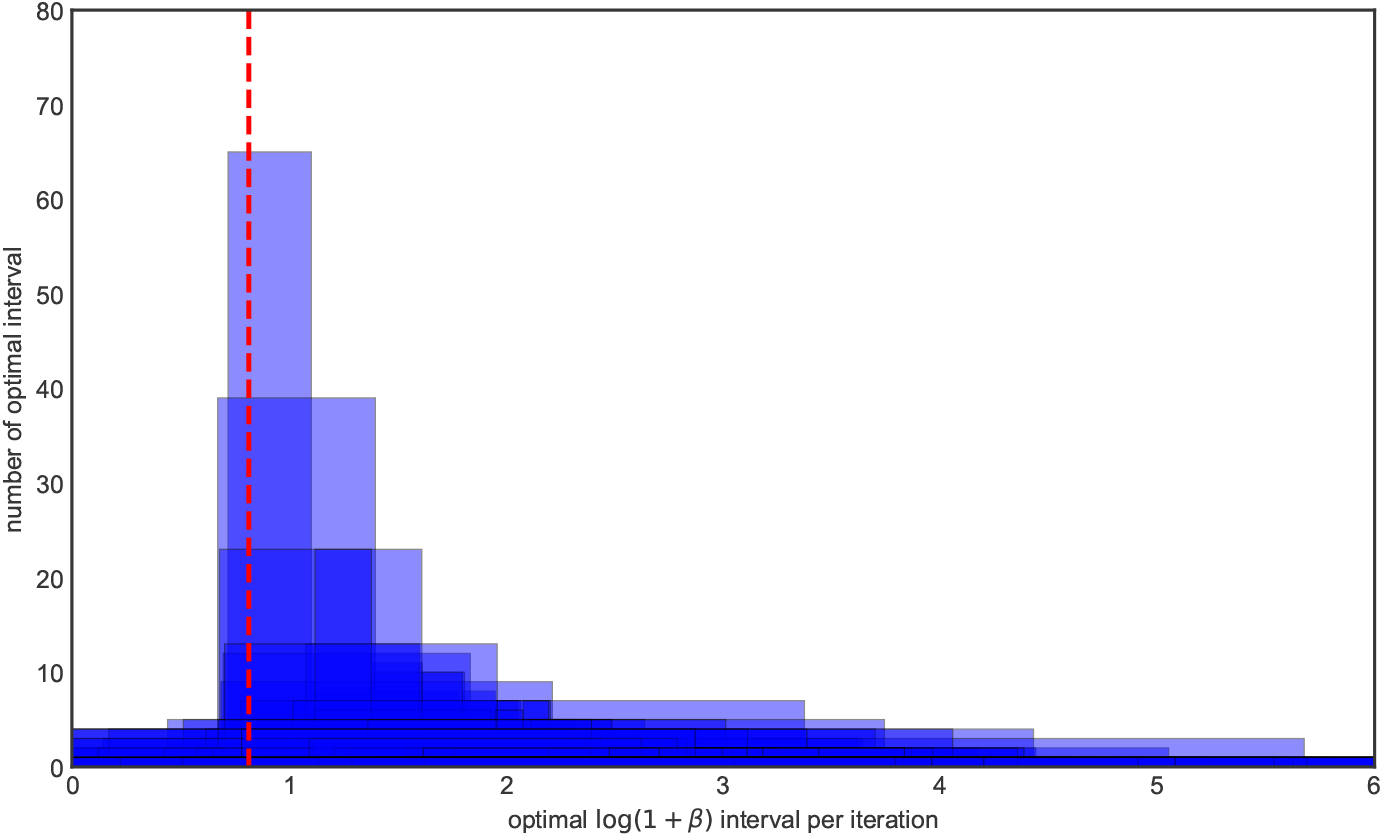
The histogram of optimal *β* intervals for prephased imputation under the Hamming loss, the *x* axis is the value of optimal log(1 + *β*) intervals in each iteration, and the *y* axis is the number of replicates in 1000 iterations. The *x* axis of the red dash line is the value of log(1 + *β*_0_) where *β*_0_ is the truth.

**Figure A2:**
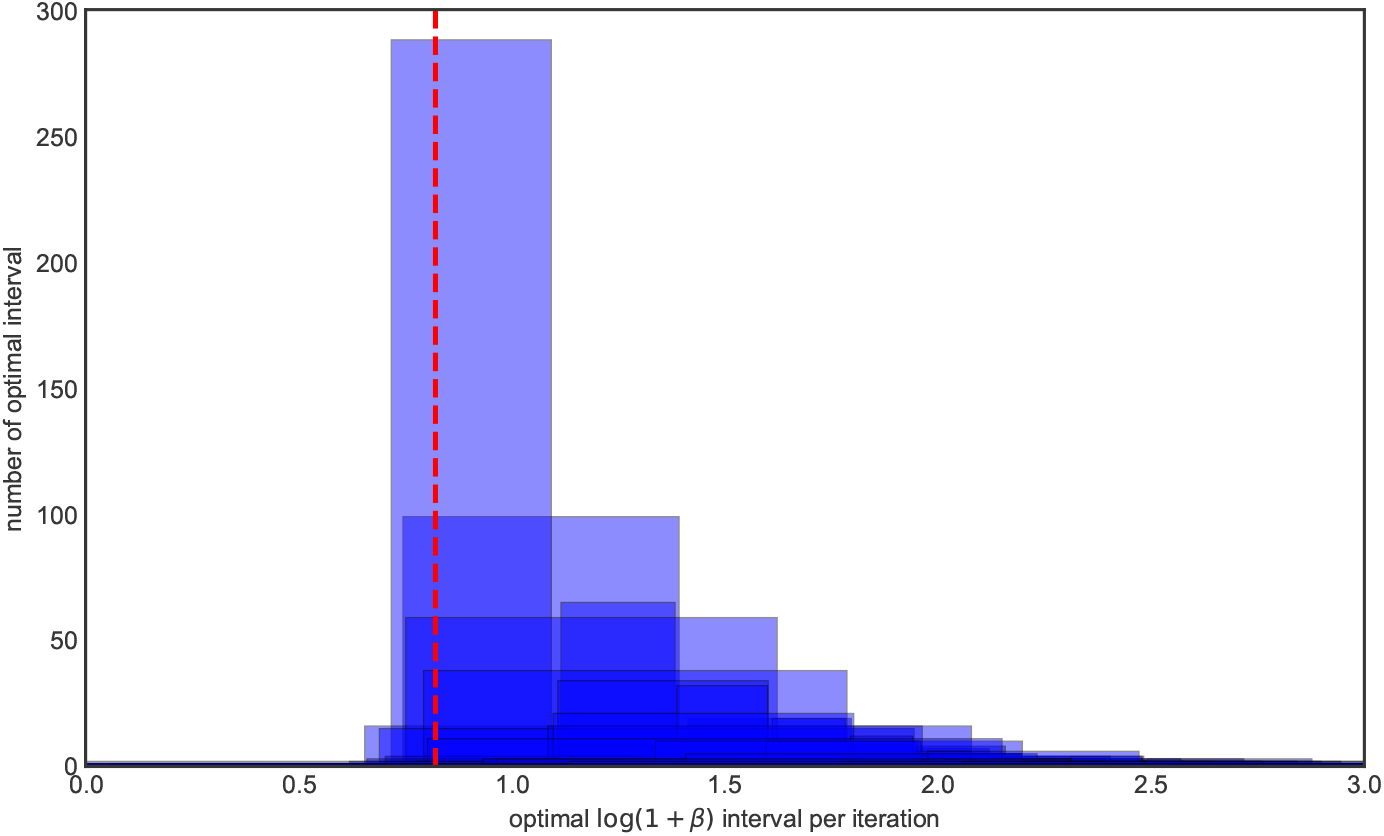
The histogram of optimal *β* intervals for prephased imputation under the Hamming loss where the model is `Africa_1T12`, the *x* axis is the value of optimal log(1 + *β*) intervals in each iteration, and the *y* axis is the number of replicates in 1000 iterations. The *x* axis of the red dash line is the value of log(1 + *β*_0_) where *β*_0_ is the truth.

**Figure A3:**
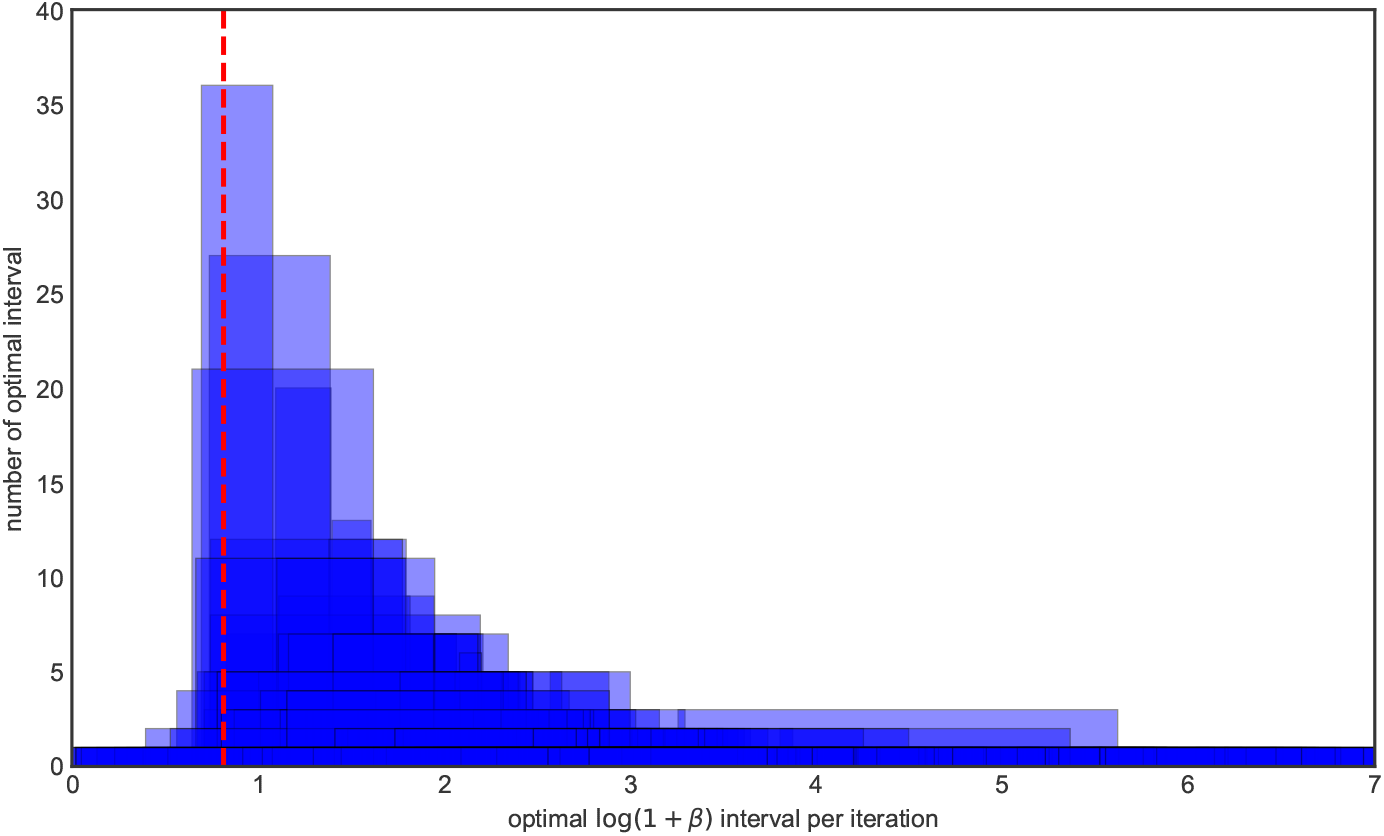
The histogram of optimal *β* intervals for prephased imputation under the variance-weighted Hamming loss, the *x* axis is the value of optimal log(1 + *β*) intervals in each iteration, and the *y* axis is the number of replicates in 1000 iterations. The *x* axis of the red dash line is the value of log(1 + *β*_0_) where *β*_0_ is the truth.

**Figure A4:**
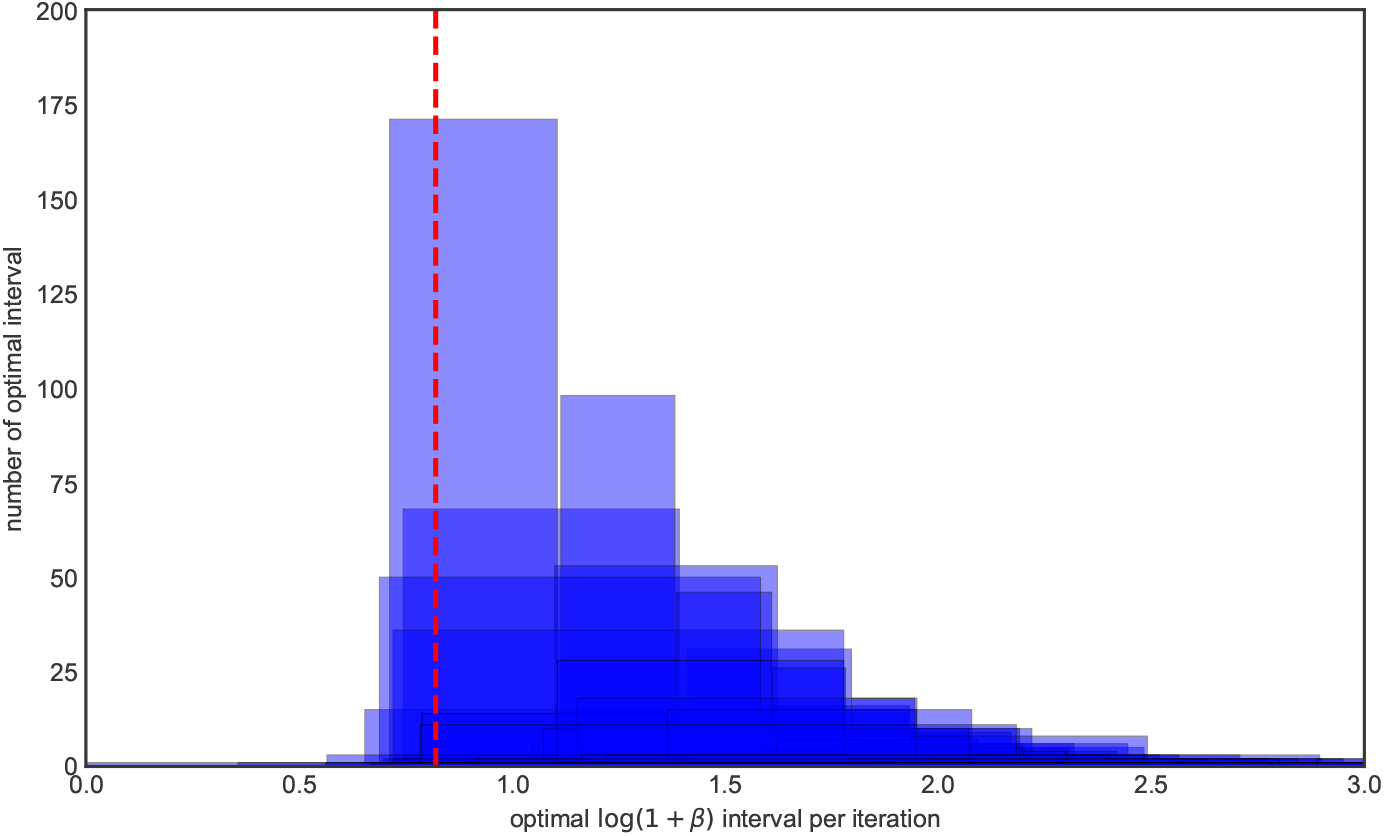
The histogram of optimal *β* intervals for prephased imputation under the variance-weighted Hamming loss where the model is `Africa_1T12`, the *x* axis is the value of optimal log(l + *β*) intervals in each iteration, and the *y* axis is the number of replicates in 1000 iterations. The *x* axis of the red dash line is the value of log(l + *β*_0_) where *β*_0_ is the truth.

## Footnotes

1 Note that, while multiple copying paths may have the same error according to equation (5), they may have different imputation errors. We verified that the results presented below were consistent from run to run, and not driven by arbitrary choices of the optimal copying path.

## References

Jeffrey R. Adrion, Christopher B. Cole, Noah Dukler, Jared G. Galloway, Ariella L. Gladstein, Graham Gower, Christopher C. Kyriazis, Aaron P. Ragsdale, Georgia Tsambos, Franz Baumdicker, Jedidiah Carlson, Reed A. Cartwright, Arun Durvasula, Bernard Y. Kim, Patrick McKenzie, Philipp W. Messer, Ekaterina Noskova, Diego Ortega-Del Vecchyo, Fernando Racimo, Travis J. Struck, Simon Gravel, Ryan N. Gutenkunst, Kirk E. Lohmeuller, Peter L. Ralph, Daniel R. Schrider, Adam Siepel, Jerome Kelleher, and Andrew D. Kern. A community-maintained standard library of population genetic models. bioRxiv, 2019. doi: 10.1101/2019.12.20.885129. URL https://www.biorxiv.org/content/early/2019/12/21/2019.12.20.885129.

A.M. Andrew. Another efficient algorithm for convex hulls in two dimensions. Information Processing Letters, 9(5):216–219, 1979. ISSN 0020-0190. doi: https://doi.org/10.1016/0020-0190(79)90072-3. URL https://www.sciencedirect.com/science/article/pii/0020019079900723.

Franz Baumdicker, Gertjan Bisschop, Daniel Goldstein, Graham Gower, Aaron P Ragsdale, Georgia Tsambos, Sha Zhu, Bjarki Eldon, E Castedo Ellerman, Jared G Galloway, Ariella L Gladstein, Gregor Gorjanc, Bing Guo, Ben Jeffery, Warren W Kretzschumar, Konrad Lohse, Michael Matschiner, Dominic Nelson, Nathaniel S Pope, Consuelo D Quinto-Cortés, Murillo F Rodrigues, Kumar Saunack, Thibaut Sellinger, Kevin Thornton, Hugo van Kemenade, Anthony W Wohns, Yan Wong, Simon Gravel, Andrew D Kern, Jere Koskela, Peter L Ralph, and Jerome Kelleher. Efficient ancestry and mutation simulation with msprime 1.0. Genetics, 220(3): iyab229, 2022.

Brian L Browning and Sharon R Browning. Genotype imputation with millions of reference samples. American journal of human genetics, 98(1):116–126, 2016. ISSN 0002-9297. doi: 10.1016/j.ajhg.2015.11.02.

Brian L Browning, Xiaowen Tian, Ying Zhou, and Sharon R Browning. Fast two-stage phasing of large-scale sequence data. The American Journal of Human Genetics, 108(10):1880–1890, 2021. ISSN 0002-9297. doi: https://doi.org/10.1016/j.ajhg.2021.08.005. URL https://www.sciencedirect.com/science/article/pii/S0002929721003049.

Petr Danecek, Adam Auton, Goncalo Abecasis, Cornelis A. Albers, Eric Banks, Mark A. DePristo, Robert E. Handsaker, Gerton Lunter, Gabor T. Marth, Stephen T. Sherry, Gilean McVean, Richard Durbin, and 1000 Genomes Project Analysis Group. The variant call format and VCFtools. Bioinformatics, 27(15):2156–2158, 06 2011. ISSN 1367-4803. doi: 10.1093/bioinformatics/btr330. URL https://doi.org/10.1093/bioinformatics/btr330.

Olivier Delaneau, Jonathan Marchini, and Jean-François Zagury. A linear complexity phasing method for thousands of genomes. Nature methods, 9(2):179–181, 2012.

Bradley Efron, Trevor Hastie, Iain Johnstone, and Robert Tibshirani. Least angle regression. The Annals of Statistics, 32(2):407–499, 2004. doi: 10.1214/009053604000000067. URL https://doi.org/10.1214/009053604000000067.

Kaylea Haynes, Paul Fearnhead, and Idris A Eckley. A computationally efficient nonparametric approach for changepoint detection. Statistics and Computing, 27(5):1293–1305, 2017. doi: 10.1007/s11222-016-9687-5. URLhttps://doi.org/10.1007/s11222-016-9687-5.

Bryan Howie, Christian Fuchsberger, Matthew Stephens, Jonathan Marchini, and Gonçalo R Abecasis. Fast and accurate genotype imputation in genome-wide association studies through pre-phasing. Nature genetics, 44(8):955–959, 2012.

Bryan N Howie, Peter Donnelly, and Jonathan Marchini. A flexible and accurate genotype imputation method for the next generation of genome-wide association studies. PLoS Genet, 5(6):e1000529, 2009.

Jerome Kelleher, Alison M Etheridge, and Gilean McVean. Efficient coalescent simulation and genealogical analysis for large sample sizes. PLoS computational biology, 12(5):e1004842, 2016.

Caleb Ki and Jonathan Terhorst. Exact decoding of the sequentially markov coalescent. bioRxiv, 2020.

Marc Lavielle. Using penalized contrasts for the change-point problem. Signal Processing, 85:1501–1510, 08 2005. doi: 10.1016/j.sigpro.2005.01.012.

N. Li and M. Stephens. Modeling linkage disequilibrium and identifying recombination hotspots using single-nucleotide polymorphism data. Genetics, 165:2213–2233, 2003.

Y. Li and G. R. Abecasis. Mach 1.0: Rapid haplotype reconstruction and missing genotype inference. Am. J. Hum. Genet., S79:2290, 2006.

Po-Ru Loh, Pier Francesco Palamara, and Alkes L Price. Fast and accurate long-range phasing in a uk biobank cohort. Nature genetics, 48(7):811, 2016.

Gerton Lunter. Haplotype matching in large cohorts using the li and stephens model. Bioinformatics, 35(5):798–806, 08 2019. ISSN 1367-4803. doi: 10.1093/bioinformatics/bty735. URL https://doi.org/10.1093/bioinformatics/bty735.

Jonathan Marchini, Bryan Howie, Simon R Myers, Gil McVean, and Peter Donnelly. A new multipoint method for genome-wide association studies by imputation of genotypes. Nat Genet, 39(7):906–13, 2007.

Joshua S. Paul and Yun S. Song. A principled approach to deriving approximate conditional sampling distributions in population genetics models with recombination. Genetics, 186:321–338, 2010.

P. Scheet and M. Stephens. A fast and flexible statistical model for large-scale population genotype data: Applications to inferring missing genotypes and haplotypic phase. Am. J. Hum. Genet., 78:629–644, 2006.

Yun S Song. Na li and matthew stephens on modeling linkage disequilibrium. Genetics, 203(3):1005–1006, 2016.

Jacob A Tennessen, Abigail W Bigham, Timothy D O’Connor, Wenqing Fu, Eimear E Kenny, Simon Gravel, Sean McGee, Ron Do, Xiaoming Liu, and Goo Jun. Evolution and functional impactofrare coding variation from deep sequencing of human exomes. Science, 337(6090):64–69, 2012.

